# A touch of hierarchy. Population Receptive Fields reveal fingertip integration in Brodmann areas in human primary somatosensory cortex

**DOI:** 10.1101/2021.01.15.426783

**Authors:** W. Schellekens, M. Thio, S. Badde, J. Winawer, N. Ramsey, N. Petridou

**Author notes:** **Corresponding author** W. Schellekens, Postal address: Q101.132, P.O.Box 85500, 3508 GA, Utrecht, Netherlands.

## Abstract

Several neuroimaging studies have shown the somatotopy of body part representations in primary somatosensory cortex (S1), but the functional hierarchy of distinct subregions in human S1 has not been adequately addressed. The current study investigates the functional hierarchy of cyto-architectonically distinct regions, Brodmann areas BA3, BA1, and BA2, in human S1. During functional MRI experiments, we presented participants with vibrotactile stimulation of the fingertips at 3 different vibration frequencies. Using population Receptive Field (pRF) modeling of the fMRI BOLD activity, we identified the hand region in S1 and the somatotopy of the fingertips. For each voxel, the pRF center indicates the finger that most effectively drives the BOLD signal, and the pRF size measures the spatial somatic pooling of fingertips. We find a systematic relationship of pRF sizes from lower-order areas to higher-order areas. Specifically, we found that pRF sizes are smallest in BA3, increase slightly towards BA1, and are largest in BA2, paralleling the increase in visual receptive field size as one ascends the visual hierarchy. Additionally, we find that the time-to-peak of the hemodynamic response in BA3 is roughly 0.5s earlier compared to BA1 and BA2, further supporting the notion of a functional hierarchy of subregions in S1. These results were obtained during stimulation of different mechanoreceptors, suggesting that different afferent fibers leading up to S1 feed into the same cortical hierarchy.

## Introduction

Touch is an important source of information on our direct surroundings. We use touch information to explore objects and surfaces and touch plays a major part in haptic processes such as tool use. The loss of adequate touch signal processing, e.g. due to stroke, frequently leads to severe impairments affecting many facets of everyday life. Hence, understanding somatosensory processes in the human brain following cutaneous touch signals is relevant to many scientific areas ranging from fundamental neuroscience to the deciphering of neurological disorders of the somatosensory system. Imaging studies in humans have mostly addressed the somatotopic organization of the hand and fingers (Maldjian et al. 1999; Kurth et al. 2000; Hlustík et al. 2001; Blankenburg et al. 2003; Nelson and Chen 2008; Schweizer et al. 2008; Sanchez-Panchuelo et al. 2010; Ann Stringer et al. 2014; Martuzzi et al. 2014; Choi et al. 2016; Kikkert et al. 2016; Kolasinski et al. 2016; Sanchez Panchuelo et al. 2018; Da Rocha Amaral et al. 2019; Puckett et al. 2020), and the whole body (Akselrod et al. 2017; Tal et al. 2017). However, other functional characteristics of human S1 have not received equal attention. Specifically, the processing hierarchy of cyto-architectonically distinct regions in human S1, i.e. Brodmann areas BA3a/b, BA1, and BA2, (Brodmann 1909; Geyer et al 1999), has been investigated structurally in humans (Sánchez-Panchuelo et al. 2014; Wagstyl et al. 2015), but not from a functional perspective. In the current study, we investigate the functional hierarchy in human S1 by estimating the integration of somatic information in different Brodmann areas

When cortical information is processed at different hierarchical levels, information from multiple lower-level sources is integrated at the higher-order level. As a result, regions of higher hierarchical order contain neurons that exhibit larger or more complex receptive fields, meaning that neurons are responsive to more input or specific combinations of input. Functional hierarchy among separate S1 regions in humans can, therefore, potentially be revealed through a form of spatial somatosensory information integration (Hubel and Wiesel 1968; Duffy and Burchfiel 1971; Van Essen and Maunsell 1983). Previous animal studies have reported that BA3b is the primary target of thalamic output from the ventrolateral and ventroposterior nucleus (Jones and Powell 1970; Chung et al. 1986; Miller et al. 2001), which then projects onwards to BA1 and BA2 (Friedman 1983; Felleman and Van Essen 1991; Kaas 1993; Iwamura 1998). As a result, neuronal receptive fields, as reported in animal studies, are smallest in BA3b and increase in size in BA1, BA2 and beyond (Armstrong-James 1975; Hyvärinen and Poranan 1978; Sur et al. 1980; DiCarlo et al. 1998). In humans, receptive field properties of individual neurons cannot easily be assessed in healthy volunteers under normal circumstances. However, average receptive field properties of small neuronal populations (e.g. neurons inside a single MRI-voxel) can be estimated using a Gaussian population Receptive Field (pRF) model. PRF modeling was originally developed for vision (Dumoulin and Wandell 2008), where it has exposed hierarchical processing characteristics as well as other traits of the human visual system (Harvey and Dumoulin 2011; Haak et al. 2012; Dumoulin et al. 2014; Klein et al. 2014; Wandell and Winawer 2015; Merkel et al. 2018; Welbourne et al. 2018). Furthermore, two recent functional MRI (fMRI) studies have shown that pRF modeling can also be used to describe the average receptive field properties of small neuronal populations in human S1 (Schellekens et al. 2018; Puckett et al. 2020). Even though some studies find evidence consistent with hierarchical organization of somatosensory processing in humans (Bodegård et al. 2001; Van Boven et al. 2005; Dijkerman and de Haan 2007; Kim et al. 2015; Whitehead et al. 2019), the extent of spatial integration across different Brodmann areas in human S1 is presently not well defined.

The current objective is to estimate pRF properties across Brodmann areas, following vibrotactile stimulation of the fingertips. Vibrotactile stimulation can be signaled by two distinct cutaneous mechanoreceptors: Meissner corpuscles and Pacinian corpuscles, depending on the frequency of vibration (Mountcastle et al. 1972; Bolanowski et al. 1988; Pasterkamp 1999). Meissner corpuscles typically show a peak activity for flutter frequencies (i.e. between 10 Hz and 50 Hz), while Pacinian corpuscles respond to higher frequencies with a preference around 250 Hz (Rowe 2002). Furthermore, previous studies showed that Meissner and Pacinian corpuscles signal somatosensory information through different pathways, i.e. Rapid-Adapting (RA) and Pacinian pathways (Vallbo and Johansson 1984; Gescheider et al. 2004; Harvey et al. 2013; Saal et al. 2015), which reportedly project to different regions of the thalamus (Herron and Dykes 1986; Kaas 1993). Additionally, Pacinian pathways may have more connections to BA1 than BA3b (Paul et al. 1972; Hyvärinen and Poranan 1978; Iwamura et al. 1993). Hence, the hierarchical order of somatosensory processing among Brodmann areas in S1 may be frequency-dependent or at least influenced by the supplied frequency of vibration. To investigate hierarchical differences caused by stimulated mechanoreceptor type, we supplied a vibrational stimulus to the fingertips at three different frequencies: 30 Hz, 110 Hz, and 190 Hz. A perfect isolation of stimulated mechanoreceptor type is not realistic and multiple pathways likely contribute to the observed cortical signal with increasing contributions of Pacinian pathways for higher stimulation frequencies (Choi et al. 2016; Kuroki et al. 2017). Thus, differences in initial cortical projection site between RA an Pacinian pathways could be detected through changes in pRF size for different vibrotactile stimulation frequencies

In the present study, we scrutinize the hierarchical organization of S1 by measuring the properties of tactile pRFs in BA3b (from here on referred to as BA3), BA1, and BA2. The five fingers of the right hand were vibrotactually stimulated at three different frequencies, 30 Hz, 110 Hz, and 190 Hz, while Blood-Oxygen-Level-Dependent (BOLD) activity in S1 was measured with 7T fMRI. PRF modeling allows us to infer the somatotopic tuning of neuronal populations in each of the three Brodmann areas. We expect an increase in pRF size, the specificity of the somatotopic tuning, along the somatosensory processing pathway. Such a finding would indicate increasing spatial integration and be in accordance with sequential information processing and increasing processing complexity from BA3 to BA1, and finally BA2. The hierarchical order across Brodmann areas is further investigated by examining the temporal dynamics of the hemodynamic response function (HRF). Finally, the effect of mechanoreceptor pathway on cortical pRF size is presently unknown. Through pRF size estimations in different Brodmann areas under different vibrotactile frequency conditions, we investigate putative differences in cortical hierarchical projections related to different mechanoreceptor types

## Material & Methods

### Participants

Eight healthy volunteers (age range 23-31 years old, 4 female) participated in the study. All participants gave written informed consent before entering the study. The protocol was approved by the local medical ethics committee of the University Medical Center Utrecht, Netherlands, in accordance with the Declaration of Helsinki (2013).

### Apparatus

The vibrotactile stimulus was delivered using MR-compatible piezoelectric stimulators with a triangular shaped tip and a contact area of approximately 1 mm^2^ (http://dancerdesign.co.uk/). The stimulation was controlled via a custom-written MATLAB (www.mathworks.com) script. Analog stimulus signals were transferred to the stimulators using a NI-9264 digital-to-analog converter output module (National Instruments, Austin, TX, USA), which was connected to a conventional laptop and an amplifier

We mounted 5 stimulators on a plexiglass plate using ordinary adhesive gum. The adhesive gum allowed for the repositioning of the 5 stimulators to match each participant’s hand. The fingertips of the right hand were placed on the stimulators (digits did not touch each other). The hand and fingers were taped to the plexiglass plate with standard paper tape to prevent the fingers from accidentally disconnecting from the stimulators. The plexiglass plate rested on the participant’s abdomen, while the right elbow was supported by towels. Using this setup, the subject could maintain a stationary position of the right arm/hand comfortably for the full length of the fMRI experiments. This minimized movement of the hands, which could affect the results. Moreover, subjects were explicitly instructed to keep both hands still during the experiments

### Procedure and stimuli

Each subject underwent 4 fMRI experiments: the first 3 were pRF experiments, conducted to estimate pRF properties (i.e. receptive field center, size, and amplitude). These 3 experiments differed only with respect to the frequency of vibration (30 Hz, 110 Hz, and 190 Hz). The 4^th^ fMRI experiment was conducted to estimate the hemodynamic response function (HRF) within each individual subject’s S1. During the 3 pRF experiments, each fingertip was stimulated 8 times in a pseudo-randomized order. Only one fingertip was stimulated at a time, and a single stimulation lasted for 4s. An intermittent stimulation paradigm was chosen to minimize adaptation processes and, therefore, maximize the observed BOLD response: during the 4s stimulation period, a 400ms on period was alternated with a 100ms off period. After the 4s stimulation period, a 10s rest period ensued except for 8 randomly selected stimulation periods when the ensuing rest period was lengthened to 14.4s. Our analysis did not require a complete return to baseline, but rather allowed for the response to one stimulus to persist into the onset of the next. In total, a single pRF experiment took 595.2s. During the HRF experiment, a brief vibrotactile stimulation of 500ms at 30 Hz was applied to all 5 fingertips simultaneously. The brief 500ms stimulation was delivered intermittently: 200ms on / 100ms off / 200ms on. There were 32 500ms events throughout the HRF experiment with variable inter-stimulus interval (ISI). The minimum ISI was 3.05s, the maximum ISI was 23.97s, and the median ISI was 7.98s.The full HRF experiment took 320s

### Scan protocol

Scanning was conducted at a 7 Tesla Philips Achieva scanner (Philips, Best, Netherlands), using a volume transmit and a 32-channel receive headcoil (Nova medical, MA, USA). A multi-slice gradient echo (GE) echo-planar imaging (EPI) sequence was used for functional image acquisition with the following specifications: TR/TE: 1600/27ms, flip angle: 70°, SENSE factor: 3 in the anterior-posterior direction, field-of view (FOV) (ap,fh,rl): 209.4 × 41.6 × 165.0 mm at 1.6 × 1.6 × 1.6 mm voxel resolution, and interleaved slice acquisition. The FOV was placed on the superior part of the brain, covering the hand region of the postcentral gyrus. 372 volumes were acquired per pRF experiment and 200 volumes were acquired for the HRF experiment. Additionally, 10 volumes were acquired with a reversed phase encoding direction (i.e. posterior to anterior) for correction of geometrical distortions. Finally, a whole-brain T1-weighted volume was acquired with TR/TE: 7.00/3.05ms, flip angle: 8°, FOV (ap,fh,rl): 250 × 200 × 190 mm at 0.78 × 0.78 × 0.8 mm voxel size, and a whole-brain proton density volume of equal dimensions

### Image processing

The T1-weighted anatomical volume was adjusted for proton density to correct for large scale intensity inhomogeneities (Van de Moortele et al. 2009). Afterwards white matter and pial brain surfaces were estimated using Freesurfer (https://surfer.nmr.mgh.harvard.edu/). These surfaces were also inflated and flattened using Freesurfer. The functional volumes were slice time corrected, realigned (i.e. corrected for head motion), corrected for geometrical distortions, and co-registered to the anatomical T1-weighted volume using AFNI. Transformation matrices for these steps were computed using the AFNI functions 3dvolreg, 3dQwarp, and 3dAllineate, respectively. The transformation matrices were combined and all spatial preprocessing transformations were applied within a single interpolation step using the AFNI function 3dNwarpApply to minimize smoothing caused by multiple interpolation steps and general interpolation errors. The functional volumes were mapped onto the estimated cortical surface reconstructions across the full depth of the estimated grey matter using Freesurfer, creating a timeseries per surface vertex. The timeseries were high-pass filtered with a cut-off at 0.01 Hz and rescaled to percent signal change. Finally, regions of interest were drawn on the reconstructed cortical surface, based on the Brodmann area atlas supplied by Freesurfer (Fischl et al. 2008). Region BA3 corresponded with atlas areas BA3a and BA3b (covering the rostral wall of the postcentral gyrus). Region BA1 corresponded with atlas area BA1 (covering the crown of the postcentral gyrus). Finally, region BA2 (covering the caudal wall of the postcentral gyrus) was based on atlas area BA2, but manually limited posteriorly at the base of the postcentral sulcus

### pRF analysis

Each vertex’ timeseries was fitted with a Gaussian receptive field model, which described the signal amplitude for any fingertip stimulation (1):

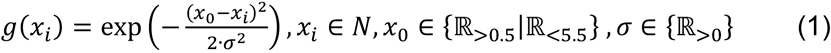

Where “*x_i_*” represents the stimulated fingertip and *“N”* is the list of fingertips ranging from 1=thumb to 5=little finger. The estimated pRF center, “*x_0_*”, describes the preferred fingertip per surface vertex and can be any real number (including fractioned numbers) between 0.5 and 5.5. A surface vertex is taken to prefer: the thumb when, 0.5<“*x_0_*”<1.5, index finger when, 1.5<“*x_0_*”<2.5, middle finger when, 2.5<“*x_0_*”<3.5, ring finger when, 3.5<“*x_0_*”<4.5, and the little finger when, 4.5<“*x_0_*”<5.5. The estimated pRF size, “*σ*”, is the spread of the Gaussian in units of fingers: the larger the pRF size, the more the neuronal population responds to stimulated fingertips in addition to the preferred one. The receptive field model “*g(x_i_)*”, then, is used to construct the effective task design (2):

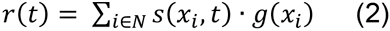

Where “*r(t)*” is the effective task design, “*s(x_i_,t)*” is the onset design matrix, which is a 2D binary matrix representing for each fingertip “*x_i_*” the stimulation onset and duration in scans *“t”*. The multiplication of the onset design matrix “*s(x_i_,t)*” and the Gaussian receptive field model “*g(x_i_)*” is summed over the fingertip dimension, resulting in the effective task design “*r(t)*”. The effective task design is convolved with a hemodynamic response function (HRF), resulting in the predicted timeseries (3):

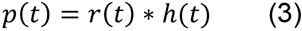

Where, “*h(t)*” is the HRF. Instead of assuming a canonical HRF, we convolved the estimated HRFs from the HRF experiment (averaged across subjects, see below) with the effective task design “*r(t)*”. Therefore, we used an HRF that was specific for each Brodmann area. The predicted timeseries model *“p(t)”* was compared with the measured timeseries of each vertex (4):

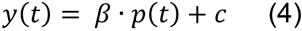

Where *y(t)* is the measured vertex’ timeseries, “*p(t)*” is the predicted timeseries, “*β*” is a scalar representing the signal amplitude and “*c*” is a constant. During the fitting procedure, optimal fits are calculated for the pRF center “*x_0_*” and size “*σ*” from equation (1) and “*β*” and “*c*” from equation (4) using the Levenberg-Marquardt (Markwardt 2009) least-square minimization algorithm (Figure 1). Finally, goodness-of-fit F-statistics were calculated for each surface vertex model fit

**Figure 1.**
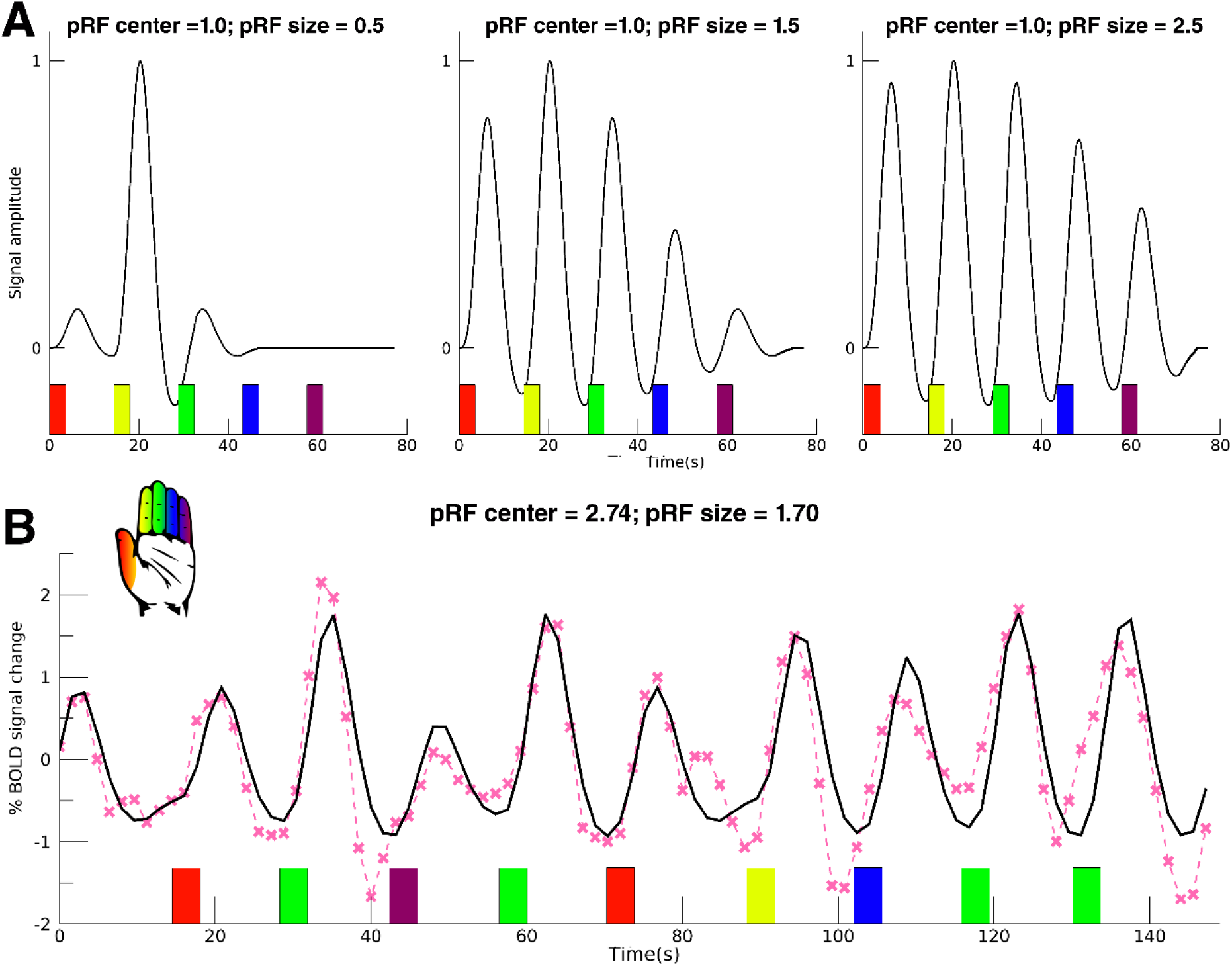
pRF model timeseries. (A) Figure shows the effect of increasing pRF size on modeled timeseries. Left image shows model with pRF center = 1 (index finger, yellow bar), pRF size = 0.5 (finger units). Middle image: pRF center = 1, pRF size = 1.5. Right image: pRF center = 1, pRF size = 2.5. The model timeseries are convolved with the average HRF from the HRF experiment and the colored bars denote the model onset time for each of the fingertip conditions, see hand icon (A) Fitted pRF timeseries (black) for one example vertex and the corresponding acquired fMRI timeseries (pink) are shown. For visibility, only a part of the complete timeseries is shown. The onsets of the fingertip stimulation conditions are represented by the colored bars, see also hand icon. This particular vertex was acquired from subject 4, BA1, 190 Hz, and was fitted with a model with pRF center= 2.74 (between index and middle finger) and pRF size = 1.70 finger units.

### HRF analysis

For the HRF experiment, we estimated the hemodynamic response function of each vertex using a set of finite impulse response (FIR) functions (Lindquist et al. 2009). The timeseries were upsampled by a factor of 4 using a 3th degree B-spline interpolation, resulting in a time point every 400ms. This matched the stimulus onset resolution, as stimulus onsets were locked to time samples every 400ms. A set of finite impulses were constructed to cover the range of 14.4 seconds (i.e. 36 finite impulses), starting from the moment of stimulation. The amplitude in percent signal change at each time point was calculated using a multiple linear regression. An HRF per ROI was created by averaging the estimated HRFs of all vertices within the ROIs that showed a significant fit with respect to the HRF task design (false-discovery-rate corrected). Afterwards, the peak amplitude, time to peak (TTP) and full-width-at-half-maximum (FWHM) were extracted from the estimated HRF curves

### Statistical analyses

For the statistical analyses of all experiments we included the surface vertices with a significant goodness-of-fit F-statistic derived from the pRF experiments (false-discovery-rate corrected) that fell in one of the three predefined ROIs. The percentage explained variance per vertex was calculated through the Pearson correlation coefficient of predicted timeseries and obtained timeseries squared. The presence of a somatotopy was assessed using the vertex coordinates of the flattened surfaces. Initially, the flattened surfaces were manually rotated so that the central sulcus was vertically aligned along the dorsoventral axis. A somatotopy is defined here as the linear relationship between dorsoventral coordinates and pRF centers. Hence, the slope between coordinates and pRF centers reflects the presence of a somatotopy and was calculated using a linear regression per ROI, per vibrotactile frequency and per subject. We used Student’s t-test to test if slopes deviated significantly from zero. We used a 2-way univariate repeated measures ANOVA with the slopes as dependent variable and ROI and vibrotactile frequency as repeated measures factors (3 levels each) to test for differences in somatotopic structures per ROI or frequency of vibration. The pRF sizes were binned in 5 preferred finger representation bins, according to the pRF centers. Then, we applied a 3-way univariate repeated measures ANOVA to test for differences in pRF size across ROI, vibrotactile frequency, and preferred finger representation (with 3, 3, and 5 levels, respectively) with linear contrasts for each factor. The same 3-way univariate repeated measures ANOVA was performed on the estimated amplitude of the percent BOLD signal change (i.e. “*β”* from equation (4)). For the HRF experiment, differences in peak amplitude, TTP, and FWHM per ROI were also tested for using univariate repeated measures ANOVAs with only ROI as factor (3 levels)

## Results

### S1 Somatotopy – spatial organization of pRFs

We used a Gaussian receptive field model to estimate the timeseries of the pRF experiments (Figure 1B). The predicted timeseries explained on average 35% (s.d.=11%) of variance of the recorded BOLD fMRI signal within the 3 predefined ROIs. On the basis of the estimated pRF centers we found the somatotopy of the five fingertips along the ventrolateral to mediodorsal axis of the postcentral gyrus in all 3 Brodmann areas (Figure 2): BA3: t_(7)_=13.10, p<0.001, BA1: t_(7)_=13.25, p<0.001, BA2: t_(7)_=8.51, p<0.001. The somatotopy, characterized as the slope of cortical coordinates and pRF centers, differed significantly across the 3 Brodmann areas (F_(2,14)_=15.26, p<0.001). Particularly, the somatotopy was less clear in Brodmann area BA2 (post-hoc somatotopy slope t-tests BA3-BA1: t_(7)_=0.55, p=0.589; BA3-BA2: t_(7)_=5.04, p<0.001; BA1-BA2: t_(7)_=4.48, p=0.001). In BA2 there appears to be a cluster of pRF centers for the thumb and index finger and a second cluster for the middle, ring and little fingers (Figure 2B). The frequency of vibration, however, did not influence the somatotopy slope (F_(2,14)_=0.25, p=0.782), although the projected somatotopy appeared less clear in several participants during the 30 Hz vibration condition compared to higher frequencies (Figure 3). We, finally, did not observe an interaction effect between Brodmann areas and applied frequency of vibration on the somatotopy slope (F_(4,28)_=0.85, p=0.505), meaning that we did not find evidence for a somatotopy change in any Brodmann area for higher frequencies

**Figure 2.**
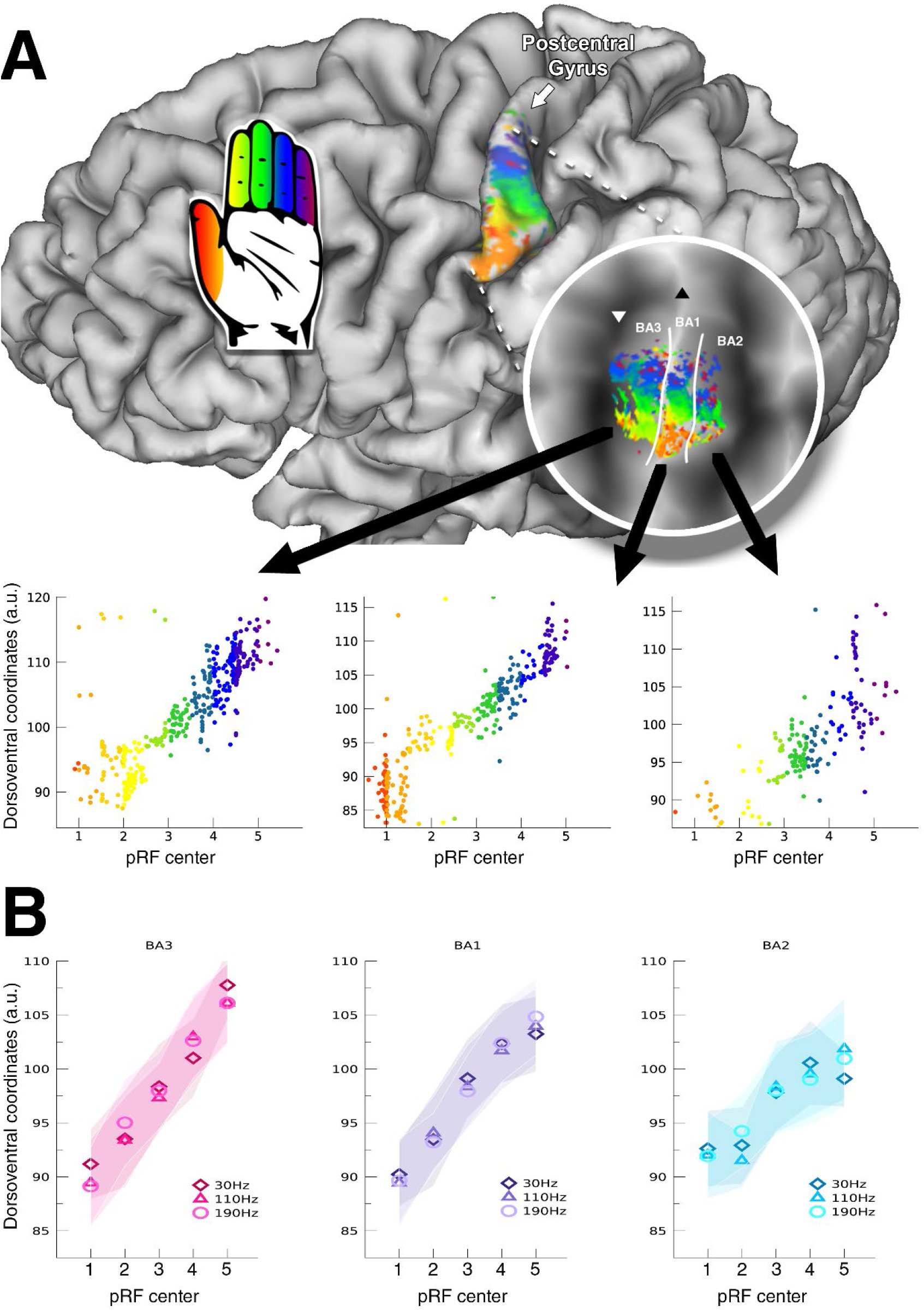
Fingertip somatotopy. (A) Single subject pRF centers following 190 Hz vibrotactile stimulation are presented on a pial surface and flattened surface (circle). The cortical coordinates along the dorsoventral axis plotted against the pRF centers are shown for all three Brodmann areas. For the pRF centers, 1=thumb, 2=index finger, 3=middle finger, 4=ring finger, 5=little finger, which is also indicated by the colors in the scatterplot and the hand icon. (B) Group average of cortical coordinates along the dorsoventral axis plotted against the mean pRF center per fingertip 1=thumb, 2=index finger, 3=middle finger, 4=ring finger, 5=little finger). Shaded area represents standard error of the mean across subjects. Different symbols represent different vibrational frequencies

**Figure 3.**
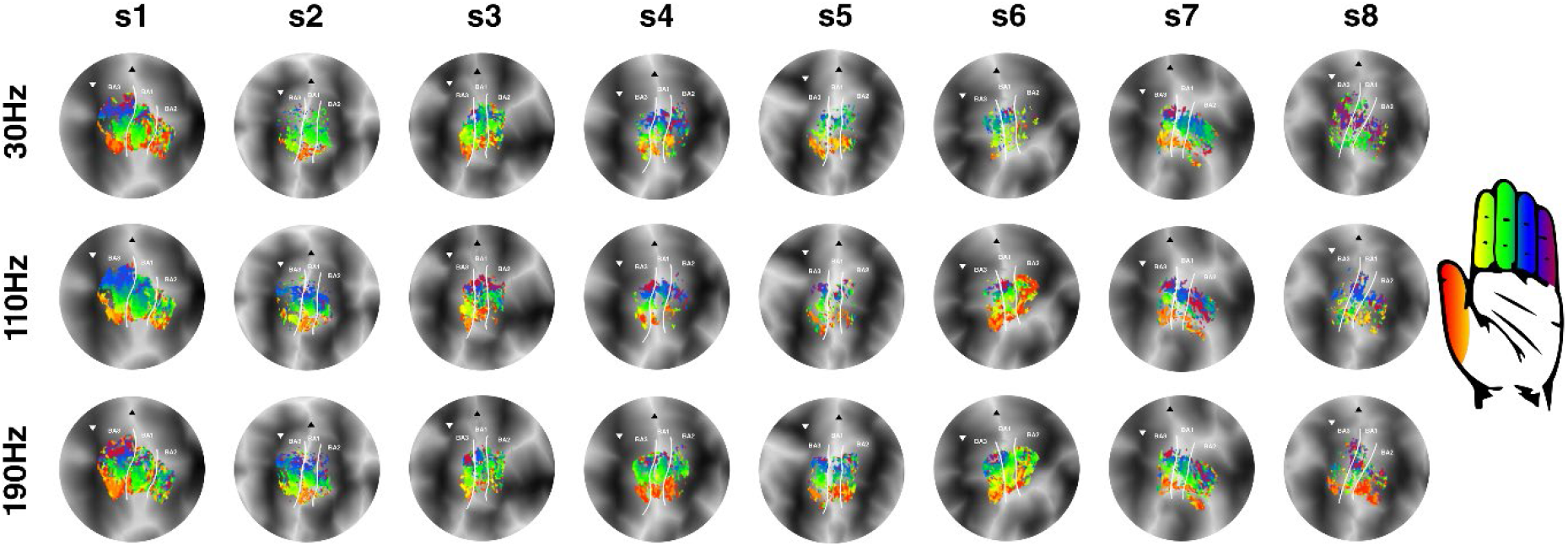
pRF center maps. The pRF centers are displayed on flattened cortical surfaces for all subjects (s1-s8). Rows depict the different frequencies of vibrotactile stimulation (30 Hz, 110 Hz & 190 Hz). Borders between Brodmann areas are denoted by the white solid line. The base of the central sulcus is shown by the white downward triangle, and the crown of the postcentral gyrus is indicated by the black upward triangle. Correspondence of pRF center and fingertip is denoted by the hand icon

### pRF sizes – fingertip specificity of the pRFs

The estimated pRF sizes (Figure 4) differed significantly across Brodmann areas (F_(2.14)_=13.26, p<0.001), showing a significant linear increase (t_(14)_=4.90, p<0.001) from BA3 to BA1 and finally BA2 (Figure 5A). The frequency of vibrotactile stimulation also influenced the receptive field sizes (F_(2,14)_=6.03, p=0.013, figure 5B), revealing a linear increase in receptive field size with an increasing vibrational frequency (t_(14)_=3.24, p=0.006). However, there was no interaction effect of frequency of vibrotactile stimulation on the included Brodmann areas (F_(4,28)_=0.69, p=0.606). Thus, we did not observe that receptive field sizes differed in any particular Brodmann area under differing vibrational frequency conditions Lastly, pRF sizes also differed per preferred fingertip (F_(4,28)_=6.90, p<0.001), which also exhibited a significant linear relationship between fingertip representation and pRF size (t_(28)_=5.13, p<0.001). Thus, pRF sizes were observed to be smallest for thumb representations and gradually increased for cortical representations of the remaining 4 fingertips, with the largest receptive field sizes for the little fingertip representations (Figure 5C). This effect of fingertip representation on pRF size did not differ among Brodmann areas (F_(8,56)_=1.32, p=0.253), or during the different frequencies of vibrotactile stimulation conditions (F_(8,56)_=1.40, p=0.217).

**Figure 4.**
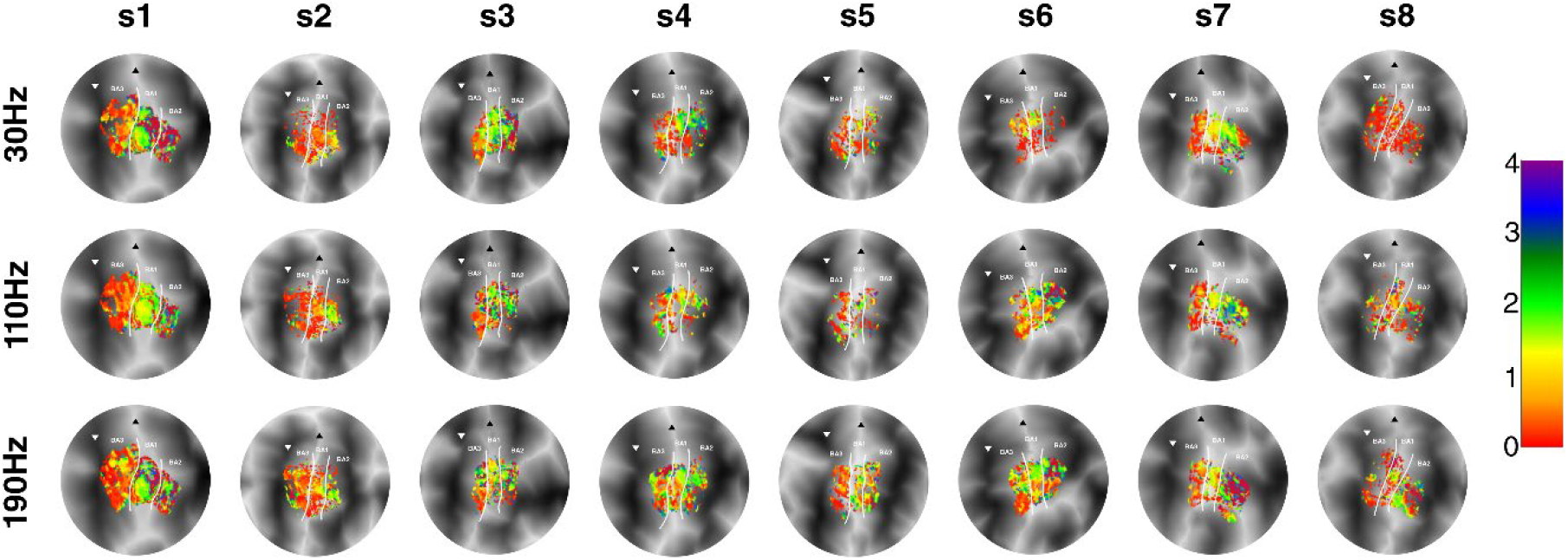
pRF size maps. The pRF sizes are displayed on flattened cortical surfaces for all subjects (s1-s8). Rows depict the different frequencies of vibrotactile stimulation (30 Hz, 110 Hz & 190 Hz). Borders between Brodmann areas are denoted by the white solid line. The base of the central sulcus is shown by the white downward triangle, and the crown of the postcentral gyrus is indicated by the black upward triangle

**Figure 5.**
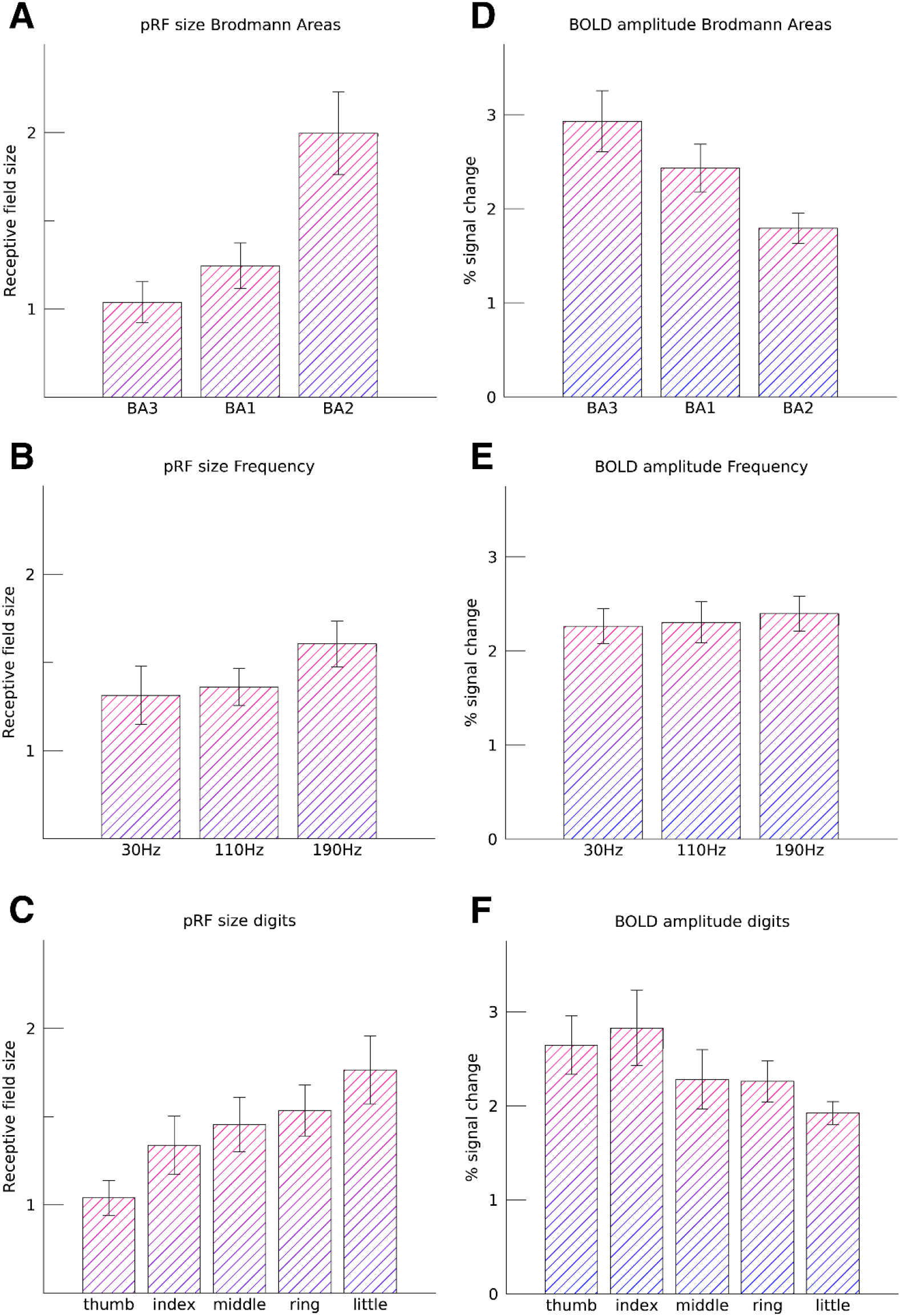
Average pRF sizes and BOLD amplitudes. Figure shows the average pRF size across subjects for Brodmann areas (A), fingertip representation (B), and vibrotactile frequency (C), as well as the corresponding estimated BOLD signal amplitude (D-F). Error bars denote the standard error of the mean across subjects

### Amplitude of the BOLD signal

We found that the amplitude of the estimated percentage of BOLD signal change (*“β”*) differed significantly across the 3 Brodmann areas (F_(2,14)_=8.15, p=0.004), where largest percent signal changes were measured in BA3 and gradually decreased towards BA2 (t_(14)_=-4.03, p=0.001, figure 5D). However, both preferred fingertip and vibrotactile frequency did not have a significant effect on the BOLD signal amplitudes (F_(4,28)_=2.21, p=0.094, and F_(2,14)_=1.75, p=0.208, respectively, Figure 5E-F). Thus, the percent BOLD signal change differed per Brodmann area, but was not significantly affected by the preferred fingertip of included populations, or by the vibrotactile frequency at which fingertips were stimulated

### Hemodynamic response function

We estimated the hemodynamic response function within S1 (figure 6). Although the largest percent signal change was observed for BA1, the peak amplitude did not deviate significantly across Brodmann areas (F_(2,12)_=2.68, p=0.109). Neither did the FWHM of the HRFs differ significantly between BA3, BA1, & BA2 (F_(2,12)_=0.97, p=0.407). However, the TTP differed significantly per Brodmann area (F_(2,12)_=5.42, p=0.021), where the TTP in BA3 was on average 0.51s (s.e.=0.17s) faster compared to the TTP seen in the other 2 Brodmann areas (post-hoc t-test BA3 – BA1+BA2: t_(12)_=3.07, p=0.010)

**Figure 6.**
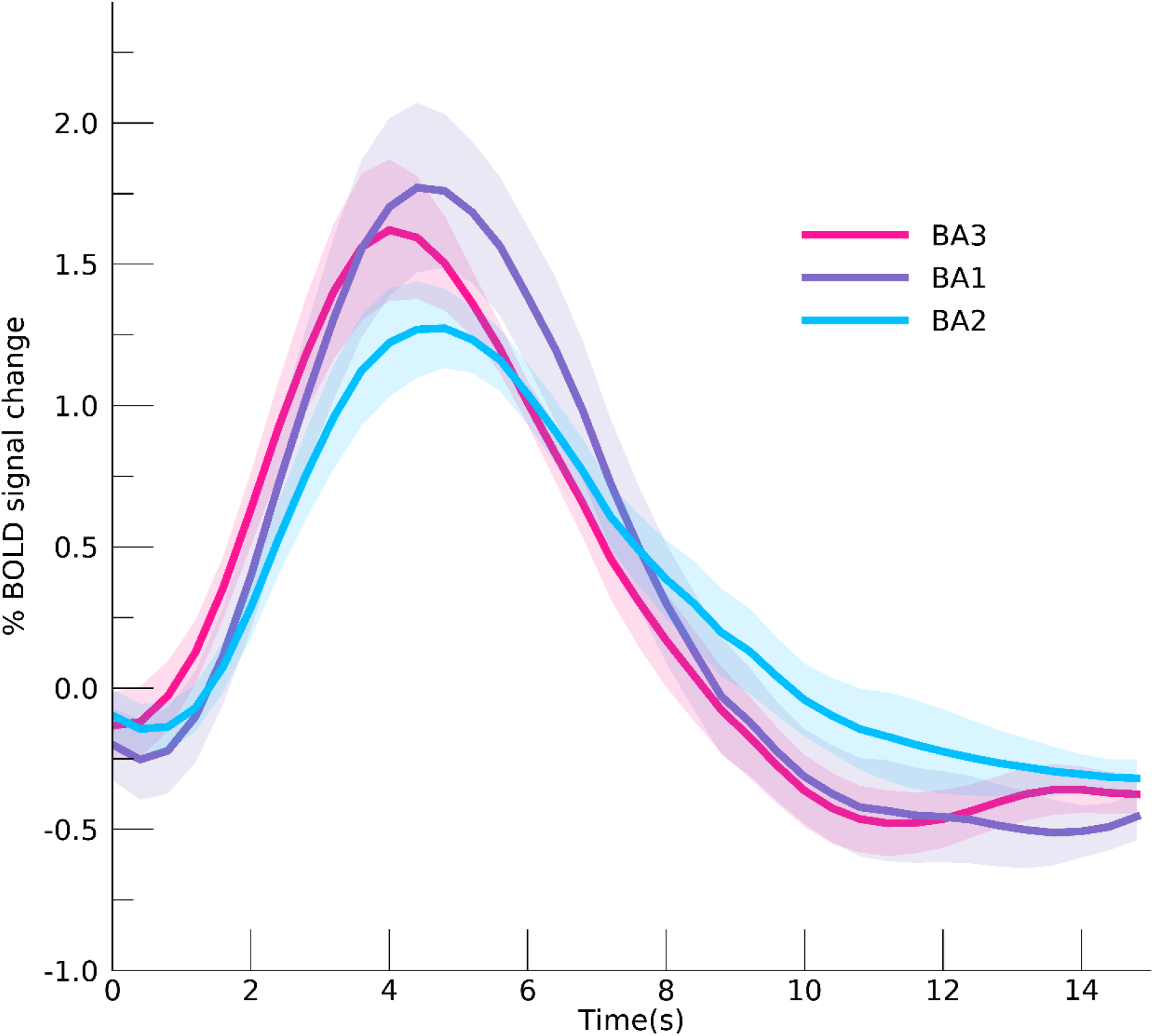
Hemodynamic response functions. Estimated hemodynamic response functions per Brodmann area. The areas denote one standard error of the mean across subjects

## Discussion

### General discussion

In the current study we estimated pRFs in 3 subdivisions of human S1. The patterns of pRFs can be used to suggest a cortical hierarchy among these areas, if we operationalize the notion of hierarchy by the size of receptive field, specifically assuming that an area with smaller pRFs is earlier in the hierarchy. We fitted a pRF model to fMRI BOLD activity in S1, following vibrotactile stimulation of the fingertips. Additionally, we stimulated at 3 different frequencies of vibration to investigate changes in pRF size across S1 related to mechanoreceptor type and corresponding afferents. We found that pRF sizes increased from BA3 to BA1 and finally BA2, consistent with the notion of a cortical hierarchy in which spatial somatic information is pooled into larger and larger regions. This effect was observed under all vibrotactile frequency conditions. PRF sizes also increased with higher frequency of stimulation. These latter two results suggests that RA and Pacinian channels share a similar cortical hierarchy, but that somatic information from a relatively larger area of the hand is pooled in S1 neuronal populations during stimulation at higher frequencies. During all frequencies of vibrotactile stimulation we observed a somatotopy of fingertips, despite the somatotopy being less clear in BA2 compared to BA3 and BA1. No significant effect of frequency on somatotopy was observed, indicating that the whole of S1 responds to vibrotactile fingertip stimulation regardless of stimulation frequency. Finally, we found that pRF sizes gradually increased from thumb to little finger. Neuronal populations that preferentially code for the thumb responded least to stimulation of other digits, compared to neuronal populations coding for the little finger, which responded to stimulation of most other digits

### Cortical hierarchy S1

Cortical hierarchy was defined in this study through information integration, which increases when information progresses higher up the processing hierarchy. Information integration is associated with the widening of response profiles of neuronal populations with respect to information coming from any number of possible sources. We estimated the widening of the response profiles of neuronal populations with a Gaussian shaped population receptive field model, where the spatial integration of somatosensory information is represented by the pRF size. We find that pRF sizes differ substantially between Brodmann areas, BA3, BA1, and BA2. Neuronal populations in BA3 have on average smallest pRF sizes, and the pRF sizes increase along the cortical processing hierarchy towards BA1 and are largest in BA2. PRF sizes in BA2 are approximately twice the size as the pRF sizes measured in BA3. This result is likely analogous to the pRF size increase among cortical areas in visual cortex, where the primary visual cortex (V1) predominantly receives thalamic output and exhibits smaller receptive field sizes than visual cortical areas further up the hierarchy, as measured both at the single unit level (Felleman and Van Essen 1991) and the population level with fMRI (Dumoulin and Wandell 2008; Wandell and Winawer 2015), which likely reflects the average receptive field size of the underlying ensemble of neurons

The hierarchical order of BA3, BA1 and BA2 is further supported by a shorter time-to-peak of the estimated HRF in BA3 compared to BA1 and BA2, which has also been observed in magnetoencephalography (MEG) studies (Inui et al. 2004; Suzuki et al. 2013) Thus, the order of cortical processing becomes apparent not merely through information integration, but also in the temporal domain. However, it is important to note that both feedforward and feedback neuronal processes contribute to the observed HRFs. Therefore, differences in temporal components of the HRF cannot solely be attributed to differences in sequential processing order. It is, for instance, possible that populations in BA1 and BA2 are not merely involved in somatosensory processing at a later point in time, but also for a slightly longer period of time, which would influence the observed HRF. Additionally, HRF latency can be affected by non-neural processes, such as the presence of draining veins (Lee et al. 1995). Nevertheless, the time-to-peak of the observed HRF in BA3 is roughly 0.5 seconds faster compared to the time-to-peak of the HRF in BA1 and BA2. Assuming factors such as draining veins don’t vary systematically between subareas in S1, this difference likely has a neuronal contribution. Our findings extend animal findings to humans, and are consistent with a cortical hierarchy in human S1, in which BA3 is the first cortical area to receive tactile information, which is then forwarded to BA1 and BA2

### Mechanoreceptive afferents

We applied three frequencies of vibrotactile stimulation to the fingertips to investigate the cortical hierarchy in human S1 as a result of different cutaneous mechanoreceptor afferents. The 30 Hz flutter frequency most likely activated Meissner corpuscles, whereas the higher frequencies would have resulted in increased contributions of Pacinian corpuscles (Bolanowski et al. 1988; Johnson 2001). Regardless of the stimulated mechanoreceptor, we observed somatotopic structures in all three included Brodmann areas. However, the somatotopy in BA2 was less clear than in the other two areas, which likely reflects less clear distinctions between cortical finger representations for areas higher up the cortical hierarchy, which has been reported in a previous animal study (Iwamura et al. 1983, 1993; Pons et al. 1985). We did not observe that the frequency of vibrotactile stimulation influenced the somatotopic structures of Brodmann areas, which may be in agreement with the notion of S1 neurons responding to multiple mechanoreceptor modalities (Pei et al. 2009; Abraira and Ginty 2013; Saal and Bensmaia 2014). However, previous optical imaging studies in monkeys have observed distinct columnar structures related to different types of mechanoreceptors in BA3. (Chen et al. 2001; Friedman et al. 2004). These frequency-dependent cortical columns are reportedly smaller than 400μm in size. The spatial resolution used in this study was not sufficiently high to capture these differences in cortical projection for different mechanoreceptor afferents

Our results show that pRF sizes increase with increasing frequency of vibrotactile stimulation. This effect was not found to differ across the three Brodmann areas and, therefore, we find no evidence to support the notion that different mechanoreceptors types project to S1 in different ways. The increase in pRF size for increased frequency could have been caused by several different processes. First, cutaneous mechanoreceptive units have receptive fields themselves, which could shape the feedforward information stream to S1 Mechanoreceptors in glabrous skin such as the Meissner corpuscle have relatively small receptive fields, whereas Pacinian corpuscles reportedly have receptive fields that extend beyond the range of one finger (Bell et al. 1994; Bolanowski and Pawson 2003). Second, neuronal activation thresholds could be dependent on vibrotactile frequency (Nelson et al. 2004; Simons et al. 2005; Ryun et al. 2017). Suprathreshold levels of activity for S1 neuronal populations could be attained during stimulation of cutaneous mechanoreceptors at high frequencies that would fall outside the neuronal populations’ receptive fields during stimulation at lower frequencies. Third, the increase in the observed pRF size for higher frequencies of vibrotactile stimulation might be an extra-classical receptive field effect (Friston 2005; Schwabe et al. 2006). It has been suggested that vibrotactile frequency discrimination is not solely driven by mechanoreceptive afferents (Kuroki et al. 2017; Birznieks et al. 2019). There may be an additional system for vibrotactile frequency processing, possibly involving horizontal connections (Schwark and Jones 1989) or the secondary somatosensory cortex (Nelson et al. 2004; Chung et al. 2013; Kalberlah et al 2013). Further research is needed to fully characterize S1 pRF properties as a function of frequency of vibrotactile stimulation

In contrast to pRF size, we did not find that the amplitude of the BOLD signal was significantly affected by frequency of vibrotactile stimulation despite the substantial difference in kinetic energy delivered to cutaneous mechanoreceptors. Previous studies, however, reported that the BOLD amplitude can either increase (Nelson et al. 2004; Goloshevsky et al. 2008) or decrease (Chung et al. 2013) for increasing vibrotactile frequencies of stimulation Especially when applying a vibrotactile stimulus for extended time periods, adaption processes might have a negative effect on the BOLD signal amplitude. For the current experiments we used an intermittent stimulation paradigm to minimize putative adaptation to the vibrotactile stimulus. It is possible that the current stimulation duration in combination with the intermittent stimulation paradigm equalized effects of different vibrotactile frequencies on BOLD amplitude

### Fingertip pRF size

We find that fingertip representations differ in pRF size. On average, cortical representations of the thumb exhibited the smallest pRF sizes, as we have reported previously (Schellekens et al. 2018). A gradual increase in pRF size is observed when progressing along the somatotopy, i.e. pRF size thumb < index < middle < ring < little finger. In a recent study, Puckett et al. 2020 reported larger pRF sizes in S1 for little finger representations compared to the index, middle and ring finger following a tactile stimulus, while measurements of the thumb were not included in their study. However, they did not observe a gradual change in pRF size across finger representations. The difference in results could possibly have been caused by methodological differences such as the smoothing applied in their analysis, which will generally increase pRF size estimates and increase the resemblance of pRF properties across voxels due to the Gaussian weighted average of neighboring voxels’ timeseries in Gaussian smoothing algorithms. Additionally, the usage of a separately estimated HRF in our study plausibly leads to better pRF estimations than using a canonical HRF as was done in the study of Puckett et al. 2020.

The difference in pRF size across fingertips occurred in all included Brodmann areas and under all vibrotactile frequency conditions. This makes it unlikely that the effect of fingertip representation on pRF size reflects functional hierarchical processes. Rather, the pRF size reflects the amount of integration of mechanoreceptive afferents from all fingers within single neuronal populations. Thus, the differences in pRF size per fingertip representation may be analogous to the increase in pRF size found in visual cortex for eccentricity representations, where foveal representations display smallest pRF sizes and outer eccentricities display larger pRF sizes (Smith et al. 2001; Dumoulin and Wandell 2008; Harvey and Dumoulin 2011). Assuming that neuronal populations representing the fovea might require high specificity for visual stimulus processing, a similar requirement may apply to somatosensory processing of tactile stimulation from the thumb and index finger. The thumb and index finger have the highest degree of motor acuity (Lachnit and Pieper 1990) and spatial acuity for somatosensory discrimination (Vega-Bermudez and Johnson 2001) Cortical pRF size might, additionally, relate to lower detection thresholds for thumb and index finger compared to the other digits in tactile discrimination tasks (Tamè et al. 2014). Our results indicate that neuronal populations that respond preferentially to the thumb and index finger receive relatively less mechanoreceptive input from the other fingers, compared to the cortical middle, ring and little finger representations, respectively

## Conclusions

We applied pRF modeling to investigate hierarchical information processing in S1 following vibrotactile stimulation of the five fingertips. PRF modeling allows for the assessment of a fingertip somatotopy in Brodmann areas BA3, BA1, and BA2. The pRF size portrays the degree of spatial information integration from the five fingertips within neuronal populations of cyto-architecturally distinct areas; smaller pRFs are associated with less spatial integration and earlier stages of the cortical processing hierarchy. pRF sizes were smallest in BA3, slightly increased for BA1, and approximately doubled in BA2, consistently across three different vibration frequencies. Additionally, we observed a difference in the time course of the hemodynamic response function among these Brodmann areas, with the shortest time-to-peak in BA3. Our findings confirm that the cortical hierarchy of the separate Brodmann areas in human S1 resembles the processing order observed in animal studies progressing from BA3 to BA1 and finally BA2, independent of the activated mechanoreceptors.

## Declarations

### Ethics approval

This study was approved by the local medical ethics committee

### Consent to participate

All participants gave written informed consent prior to inclusion

### Availability of data, material and code

All data can be made available

### Authors’ contribution

Conceptualization: WS, MT, SB, JW, NR, NP

Data acquisition: WS, MT

Analysis: WS, MT Writing: WS

